# Saccades, attentional orienting and disengagement: the effects of anodal tDCS over right posterior parietal cortex (PPC) and frontal eye field (FEF)

**DOI:** 10.1101/2021.01.25.428095

**Authors:** Lorenzo Diana, Patrick Pilastro, Edoardo N. Aiello, Aleksandra K. Eberhard-Moscicka, René M. Müri, Nadia Bolognini

**Affiliations:** School of Medicine and Surgery, University of Milano-Bicocca, Monza, Italy; Department of Psychology, University of Milano-Bicocca, Milan, Italy; Perception and Eye Movement Laboratory, Departments of Neurology and BioMedical Research, Bern University Hospital Inselspital, University of Bern, Switzerland; Department of Neurology, Bern University Hospital, Inselspital, University of Bern, Switzerland; Department of Psychology & Milan Center for Neuroscience (NeuroMI), University of Milano-Bicocca, Milan, Italy; Laboratory of Neuropsychology, Istituto Auxologico Italiano, IRCCS, Milan,Italy

**Keywords:** Visuo-spatial attention, tDCS, Saccades, Frontal Eye Field, Posterior Parietal Cortex

## Abstract

In the present work, we applied anodal transcranial direct current stimulation (tDCS) over the posterior parietal cortex (PPC) and frontal eye field (FEF) of the right hemisphere in healthy subjects to modulate attentional orienting and disengagement in a gap-overlap task. Both stimulations led to bilateral improvements in saccadic reaction times (SRTs), with larger effects for gap trials. However, analyses showed that the gap effect was not affected by tDCS. Importantly, we observed significant effects of baseline performance that may mediate side- and task-specific effects of brain stimulation.

## 1 Introduction

Visuo-spatial attention allows us to react, search and select relevant pieces of information in the surrounding space. Such an important aspect of cognition has been widely studied and multiple models have been conceptualized, capitalizing on numerous experiments of cognitive and clinical neuroscience [Posner and Petersen 1990; Corbetta and Shulman 2002]. In order for our attention to be allocated on a target, we first need to disengage it from a previous focus and subsequently orient it to this target [Posner and Petersen 1990]. Although attention can be oriented covertly (i.e., without moving the eyes), in most situations, eyes and attention move together to bring a specific stimulus within the attentional focus. Indeed, eye movements and attention are intimately connected as far as their neural underpinnings are concerned. Corbetta and Shulman [2002] described two networks for visual orienting comprising different frontal (e.g., frontal eye fields, FEFs) and temporo-parietal areas: a dorsal network for top-down, voluntary attention and a ventral network for bottom-up, automatic attention. At the same time, FEF has long been recognized as key region for the control of eye movements [Robinson and Fuchs 1969] and especially the generation of saccades, sending and receiving projections from cortical regions (i.e., frontal, parietal, and occipital) and subcortical structures, such as the superior colliculus and the basal ganglia [Schall 2009]. The posterior parietal cortex (PPC) is another key region for both attention and eye movements; it is a hub of multisensory processing [e.g., Bolognini et al. 2010], and part of its neural population, the so-called parietal eye field (PEF), is involved in visuo-motor activity, including saccadic planning [Ptak and Müri 2013]. It is thus meaningful to investigate attention and eye movements from a joint perspective.

Several works attempted to modulate spatial attention and eye-movements with non-invasive brain stimulation techniques such as transcranial magnetic stimulation (TMS) and transcranial direct current stimulation (tDCS). Specifically, tDCS can be used to polarize the underlying neural tissue - with effects extending to structurally and functionally connected areas – in order to increase or decrease its responsiveness in a given task/to a given stimulus [Thair et al. 2017]. Tipically, anodal tDCS is applied to increase cortical excitability (i.e., depolarization), whereas cathodal tDCS is applied to reduce it (i.e., hyperpolarization). Although this approach proved to be valid on primary cortices [Nitsche and Paulus 2000], polarization of associative cortices yielded mixed findings with respect to excitatory/inhibitory influences on behavior. With regard to eye movements, some studies applied tDCS over the FEFs in pro- and anti-saccades tasks [Kanai et al. 2012; Tseng et al. 2018; Reteig et al. 2018]. However, the results are not conclusive: whereas Kanai et al. [2012] found stimulation- and task-specific effects on saccades directed to the contralateral visual hemifield, Tseng et al. [2018] observed that effects of anodal tDCS over the right FEF depended on the probability of target location and the individual level of performance. On the other hand, Reteig et al. [2018] found no effects of either anodal or cathodal stimulation. With respect to spatial attention, most studies employed behavioral paradigms (without eye movements recording), mostly stimulating PPC [e.g., Sparing et al. 2009; Bolognini et al. 2010; Loftus and Nicholls, 2012] and only few targeted also frontal areas [Ball et al. 2013; Roy et al. 2015]. Overall, anodal stimulation of right PPC seems to improve responsiveness to targets in the contralateral, left hemifield [Sparing et al. 2009; Roy et al. 2015]. However, Ball et al. [2013], in a conjunction search task, found no effects of right anodal parietal (PPC) or frontal (FEF) tDCS. So far, no study has ever applied tDCS to the parietal cortex to modulate saccadic parameters.

In the present work, we investigate the behavioral effects of anodal tDCS applied over two key areas for eye movements and visuospatial attention, i.e., FEF and PPC of the right hemisphere. To this end, a simple saccadic task was administered before and immediately after brain stimulation. We tested attentional orienting by measuring participants’ saccadic latency – i.e., saccadic reaction times (SRTs) - in response to targets presented on either the left or the right visual hemifields. Moreover, given the central role of PPC in disengaging attention for reflexive saccades [Müri and Nyffeler 2008], we included both gap and overlap trials [Saslow 1967]. In gap trials, disengagement is facilitated by the offset of the central fixation before the onset of the target. This effect, known as *Gap Effect* (GE), was analyzed by taking into account gap and overlap trials together, thus obtaining a measure of the costs of disengagement [Paladini et al. 2016]. Specifically, we aimed to assess whether increasing activity of the right PPC and of the right FEF would differentially enhance contralateral, leftward SRTs and whether such an effect would be modulated by the level of required attentional disengagement. Finally, since the effects of brain stimulation are non-linear and largely depend on individual factors [see e.g., Benwell et al. 2015], we explored the association between tDCS effects and attentional baseline performance.

## 2 METHODS

### 2.1 Participants

Twenty healthy volunteers (7 males and 13 females, mean age= 26±3,2 years) were recruited at the University of Milano-Bicocca. Inclusion criteria for the study were: right-handedness according to the Edinburgh Handedness Inventory [Oldfield 1971], normal or corrected-to-normal visual acuity, and absence of contraindications to tDCS [Bikson et al. 2016; Thair et al. 2017]. The study was approved by the Ethics Committee of the University of Milano-Bicocca and it was conducted in accordance with the ethical standards of the Declaration of Helsinki. All participants provided their written informed consent to the experiment.

### 2.2 Gap-overlap task

Participants performed the gap-overlap task in a dark room, seated in front of a monitor (Acer HN274H 27”) at a viewing distance of 83cm, kept constant by means of a chin-and-head rest. Eye movements were recorded by an EyeLink 1000 (SR Research Ltd., Canada). At the beginning of the task, the eye tracker was calibrated using a 9-point grid and the mean gaze accuracy was kept, on average, around .5° of visual angle. In the gap–overlap task participants were asked to perform saccades from a central fixation point towards a lateral target as quickly and accurately as possible. Target appeared randomly in two positions: 10° to the right or to the left of the central fixation point. In the gap trials the fixation point disappeared 200 ms before the appearance of the target whereas in the overlap trials the fixation remained present when the target appeared. The duration of central fixation varied between 1200 ms and 1500 ms. Each trial was separated by a 1700 ms black screen. The task involved 64 trials for a total duration of 4 minutes. Thirty-two trials were gap and 32 were overlap. In half trials the target appeared to the left and in the remaining trials to the right of the central fixation. A short break was allowed in the middle of the task to avoid excessive eye fatigue.

### 2.3 tDCS protocol and experimental procedure

tDCS was delivered by a battery-driven current stimulator (BrainSTIM device, E.M.S., Bologna, Italy; http://www.emsmedical.net/) using two electrodes (target electrode: 5×5cm^2^ and reference electrode: 7×5cm^2^) inserted into saline-soaked sponges. Anodal tDCS was applied to the target areas (i.e., right FEF and right PPC) for a duration of 10 minutes at 1 mA intensity [as in Sparing et al. 2009], with 10 seconds fade-in and fade-out. In the case of sham tDCS, the stimulator was turned off after 30s. Each participant, on three different days, underwent all three experimental tDCS conditions: right FEF, right PPC and sham (for half of the subjects over the FEF and for the other half over PPC). The order of stimulations was counterbalanced. Target areas were marked on an elastic cap that was centered on participants’ head. FEF and PPC had been previously identified on the same cap by means of a neuronavigation procedure (Softaxic 2.0, E.M.S., Bologna, Italy) on 10 healthy volunteers. The stereotaxic MNI coordinates were: 44, -66, 43 for right PPC [corresponding to P4 of the 10-20 system; Koessler et al. 2009] and 23, -13, 59 for right FEF [Kincade et al. 2005], i.e., between C4 and CF4 locations of the 10-20 system. A simulation of tDCS-induced electric field was performed with SimNIBS 3.2 [Thielscher et al. 2015] and it is depicted in Figure 1. The anode was placed over the right FEF or PPC depending on the condition, whereas the cathode (i.e., the reference electrode), was always located over the left forehead, in a supraorbital position. The electrodes were secured using two elastic bands.

**Figure 1:**
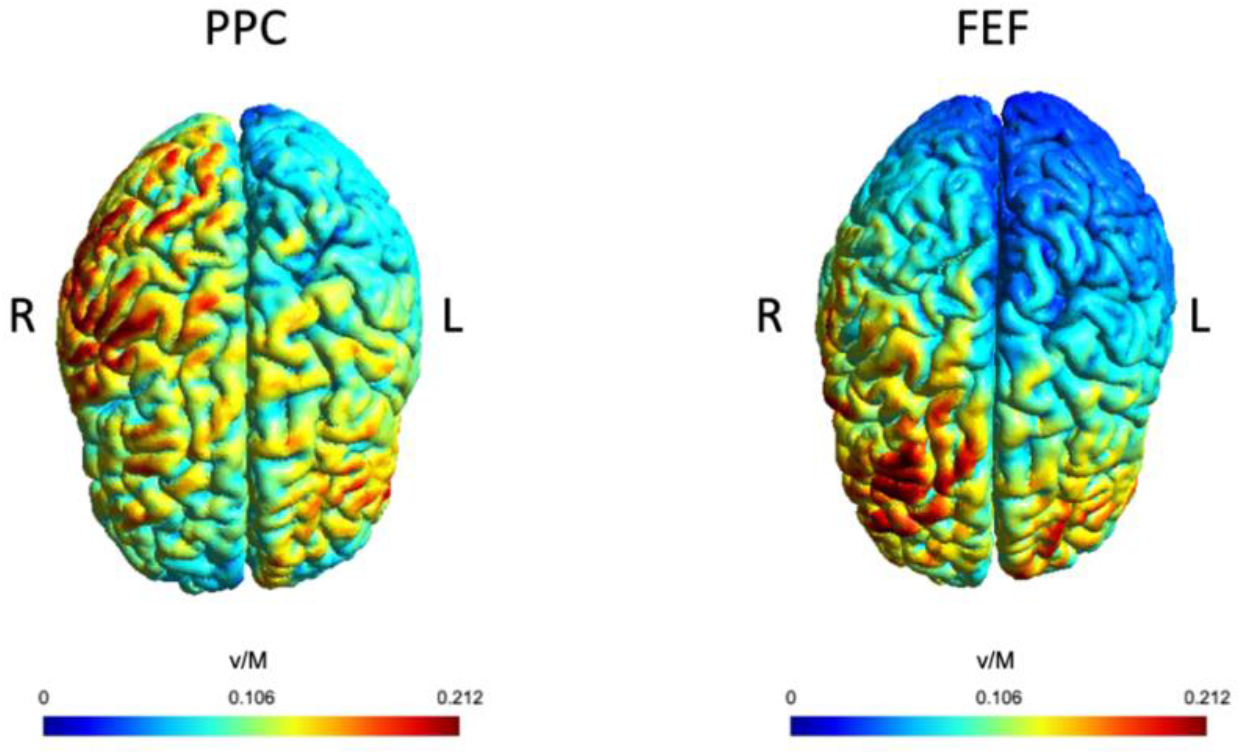
Simulated electric field (V/m) for posterior parietal cortex (PPC) and frontal eye field (FEF) anodal tDCS. R= right hemisphere; L= left hemisphere.

During each experimental session, participants performed the gap-overlap task before and right after tDCS. During the stimulation, they were asked to relax and look at a blank screen. Each session took place at the same time of the day and was separated by at least 24 hours to avoid any possible carry-over effects. At the end of the session, a questionnaire was administered to participants to collect sensations experienced during the stimulation [Fertonani et al. 2015]. At the end of the last session, participants were asked whether they received real or sham stimulation and when. tDCS was well tolerated and no serious adverse effects occurred. The most reported sensation was head itching of mild intensity, which began at stimulation onset and quickly stopped. For more details, see Section A.1 of the Appendices. With respect to blinding to stimulation, only four participants correctly identified all three stimulations.

### 2.4 Data analysis

Because of gaze calibration issues, one participant dropped out and was excluded. Therefore, data of 19 participants were analyzed. Anticipatory/express saccades (i.e., SRTs < 80 ms), multiple-step saccades, as well as saccades in the wrong direction with respect to the target were excluded [as in Paladini et al. 2016]. Overall, this procedure led to the exclusion of 3.45% of data, of which 2.78% were gap .67% were overlap. Analyses were performed on saccadic reaction times (SRTs), reflecting the saccadic latency from target appearance. Distributional assumptions checks were performed by means of descriptive and test statistics, as well as visual inspections [Ghasemi and Zahediasl 2012; Kim 2013]. α was set at .05. Data processing and plots were realized using R 3.6.2 [R Core Team 2019] and specific packages [Wickham et al. 2019] within R-Studio 1.2.5033 [RStudio Team 2019]. Analyses were performed with jamovi 1.6.9.0 [the jamovi project 2020], and prediction models implemented *via* its GAMLj module [Gallucci 2019];

#### 2.4.1 Gap-Overlap SRTs

As non-aggregated raw SRTs for were heavily right-skewed, we adopted a generalized linear mixed model (GLMM) assuming a Gamma distribution. We tested a 4-way *Stimulation* (levels: FEF, PPC, and sham)\**Time point* (levels: pre- and post-stimulation)\**Trial Type* (levels: gap and overlap)\**Side* (Left and right saccades) term regarding participants as clusters. A random *Stimulation*\**Timepoint* slope was entered to take into account the inter-individual variability of response to stimulation, but it yielded singularities. Therefore, a random intercept only was fitted. Deviation coding was adopted for factors. Bonferroni-corrected comparisons were also performed.

#### 2.4.2 Gap Effect (GE)

GE was calculated for each subject, stimulation, time-point, and side of saccade by subtracting median gap SRTs from median overlap SRTs. Therefore, bigger values (i.e., bigger differences between gap and overlap SRTs) were interpreted as higher costs of disengagement, whereas lower values as lower costs. As GE values were judged as being Normally distributed, we tested a three-way *Stimulation*\**Timepoint* \**Side* term employing a LMM. Participants were regarded as clusters and a random intercept only was fitted in the model (a random *Stimulation*\**Timepoint* term yielded singularities). Deviation coding was adopted for factors. Bonferroni-corrected post-hoc comparisons were performed.

#### 2.4.3 Effects of baseline performance

Since tDCS efects may vary across subjects depending on individual parameters such as baseline performance, we ran correlations between SRTs at baseline (i.e., prior to stimulation) and the change of SRTs before and after stimulation (i.e., Δ SRTs). Separate sets of correlations were run for gap and overlap trials. For each side of saccade and stimulation, Pearson correlations were calculated between baseline median SRTs and ΔSRTs, obtained by subtracting median SRTs prior to stimulation from median SRTs after the stimulations. Negative values represent faster post-stimulation SRTs, whereas positive ones correspond to slower post-stimulation SRTs. Bonferroni corrections for multiple comparisons were applied to each set of correlations, i.e., statistical significance was set at α = .5/6 = .008.

## 3 RESULTS

### 3.1 Gap-Overlap SRTs

The GLMM model revealed a significant main effect of *Stimulation* (χ^2^_2_=9.13; *p*=.010), *Timepoint* (χ^2^_1_=41.29; *p*<.001), *Trial Type* (χ^2^_1_=3019.28; *p*<.001), and *Side* (χ^2^_1_=5.31; *p*=.021). Of relevance, overlap trials were associated with slower SRTs (*M*=175 ms; *SE*=4.29 ms) than gap trials (*M*=231 ms; *SE*=4.3 ms). Notably, we observed a significant *Stimulation*Timepoint* interaction (χ^2^_2_=7.71; *p*=.021): SRTs were significantly faster after tDCS delivered to either FEF (ΔPre-Post, i.e., pre *minus* post tDCS= 6.59 ms; *p*=.003) and PPC (ΔPre-Post= 10.16 ms; *p*<.001), but not after sham stimulation (ΔPre-Post = 3.25 ms; *p*=.1). Finally, the interaction *Trial Type*\**Timepoint* (χ^2^_1_3.18; *p*=.039) showed that a post-stimulation reduction in SRTs was more pronounced for gap trials (ΔPre-Post = 8.81 ms; *p*<.001) than overlap trials (ΔPre-Post= 4.53 ms; *p*=.036). See Section A.2 of the Appendices for details.

### 3.2 Gap Effect (GE)

Overall, a trend for *Timepoint* (*F*_1,198_=3.56; *p*=.061) was observed. Moreover, a significant effect of *Side* (*F*_1,198_=5.17; *p*=.024) was also found, with left GE (*M*=59.3 ms; *SE*=4.5 ms) higher than right GE (*M*=52.2 ms; *SE*=4.5 ms). No other effects reached the significance level. Results are depicted in Figure 2, and further details are available in the Section A.3 of the Appendices.

**Figure 2:**
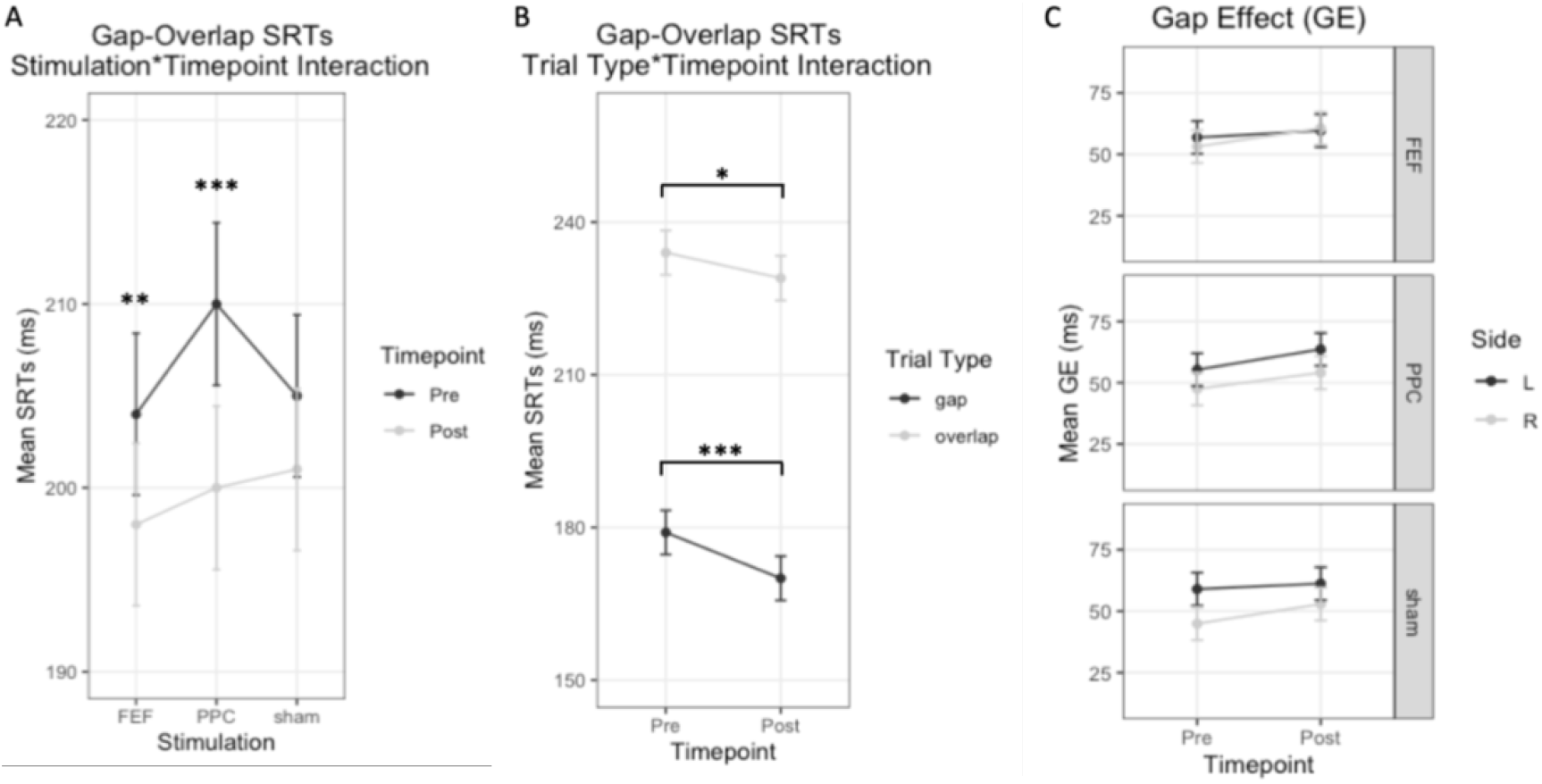
Results from the analyses of Gap-Overlap saccadic reaction times (SRTs) and gap effect (GE). A-B: mean Gap-Overlap SRTs of the significant interactions; C= mean gap effect (GE). Error bars represent the standard error. FEF= frontal eye field; PPC= posterior parietal cortex; L= left; R= right; ***= *p*<001; **= *p*<.01; *= *p*<.05.

### 3.3 Effects of baseline performance

We observed negative correlations between the baseline performance and post-pre tDCS changes, namely, participants with slower SRTs at baseline showed greater improvements after real tDCS, as indexed by more negative ΔSRTs values. Specifically, for the gap trials, we found a negative correlation for left saccades when tDCS was applied over FEF (*r*_17_=-.63; *p*=.004) and over PPC (*r*_17_=-.69; *p*=.001). As for the overlap trials, negative correlations emerged for right (*r*_17_=-.7; *p*=.001) and left (*r*_17_=-.65; *p*=.003) saccades in the FEF condition (see Table 1).

**Table 1:** Correlations between pre-stimulation saccadic reaction times (SRTs) and post-pre changes (ΔSRTs) for posterior parietal cortex (PPC), frontal eye field (FEF), and sham stimulations

| Variables | Side of Saccade | FEF | PPC | sham |
| --- | --- | --- | --- | --- |
| Gap SRTs | Left | $r_{17}=-.63; p=.004$ | $r_{17}=-.69; p=.001$ | $r_{17}=-.35; p=.14$ |
| Gap SRTs | Right | $r_{17}=-.4; p=.087$ | $r_{17}=-.4; p=.091$ | $r_{17}=-.47; p=.043$ |
| Overlap SRTs | Left | $r_{17}=-.65; p=.003$ | $r_{17}=-.48; p=.038$ | $r_{17}=-.54; p=.017$ |
| Overlap SRTs | Right | $r_{17}=-.7; p=.001$ | $r_{17}=-.11; p=.64$ | $r_{17}=-.16; p=.51$ |

## 4 DISCUSSION

In the present study we investigated the effects of anodal tDCS of right FEF and right PPC on attentional orienting in a saccadic task with different levels of disengagement. The main findings are: 1) a general enhancement of saccadic performance (i.e., faster SRTs), after both FEF and PPC stimulations, but not after sham tDCS; 2) larger enhancement effect for gap trials; 3) association between the baseline performance and tDCS effects.

### 4.1 Non-lateralized effects of FEF and PPC stimulation

The present study shows a reduction of SRTs after a single application of anodal tDCS over FEF or PPC, but not after the delivery of sham tDCS. This enhancement was not visual field-specific, i.e., it was present for both right- and left-sided saccades. With respect to FEF stimulation, unlike Kanai [2012], we did not observe a selective contralateral improvement for leftward saccades. However, our study differs in several ways from Kanai’s [2012], including tDCS montage (i.e., here a supraorbital reference), current density (i.e., here bigger electrodes), and, importantly, they delivered tDCS online, namely, during the task. It is well known that the effects of brain stimulation are state-dependent and they change according to level of activation of the brain [see e.g., Fertonani and Miniussi 2017]. Moreover, as FEF is part of the dorsal network for endogenous attentional orienting [Corbetta and Shulman 2002] and it is more involved in volitional saccades [Müri and Nyffeler 2008], we may observe effects of FEF tDCS in a task with a stronger load on top-down processes.

Concerning PPC, to our knowledge, this is the first study to apply neuromodulation over parietal areas to influence saccades. The right PPC plays a main role in the generation of exogenous saccades [Müri and Nyffeler 2008], as those in our task. Indeed, here we found that tDCS over this area brought about the largest SRTs reduction but, again, without differences between left and right saccades. We adapted the stimulation protocol from Sparing et al. [2008] who were able to improve detection for contralateral targets in a computerized task. However, unlike Sparing et al. [2008], we positioned the reference electrode more distant from the target electrode (i.e., supraorbitally) in order to allow more current reach the brain [Miranda et al. 2006], at the cost of a lower spatial focality which may explain the lack of a lateralized effect. Nonetheless, there are other possible explanations for the tDCS-induced non-lateralized, bilateral decrease of SRTs. Firstly, although a practice effect must be considered, it is possible that anodal stimulation brought about an additive effect by means of an up-regulation of right-hemispheric fronto-parietal circuits of tonic alertness, thus leading to generally faster SRTs [Sturm et al. 1999; Petersen and Posner, 2012]. This hypothesis could be explored by analyzing tDCS effects on pupil size as a proxy of arousal [Morad et al. 2000; Paladini et al. 2017]. Furthermore, our finding may also reflect what is found in lesion (or TMS “virtual lesions”) studies in humans. For example, Pierrot-Deseilligny [1991] found that brain-damaged patients with right parietal lesions showed a bilateral increase of saccadic latency. Moreover, Nyffeler et al. [2006] found a bilateral increase of SRTs after low-frequency repetitive TMS of the right FEF [see also Kapoula et al. 2001].

### 4.2 Disengagement

To study the effects of tDCS on attentional disengagement, we included in the task both gap and overlap trials. Overall, gap trials were faster than overlap ones, thus confirming the validity of the experimental paradigm [Saslow 1967]. We subsequently calculated the GE to obtain a measure of disengagement costs [Paladini et al. 2016]. Results showed that GE was not affected by anodal tDCS. We only observed a trend towards an increase of GE, irrespective of the type stimulation, which seemed primarily driven by a reduction of SRTs in gap trials, rather than by a SRTs slowdown in overlap trials. In other words, after active or sham tDCS participants showed a greater benefit of the fixation offset of the gap trials rather than an increase cost of the disengagement. Nonetheless, in the attempt to observe more subtle differences between gap and overlap trials, we run separate follow-up analyses for each trial type (See Section A.4 of the Appendices). Although these explorative analyses should be interpreted with caution, they seem to support previous results: the most pronounced reduction in SRTs for gap trials occurred after FEF and PPC tDCS, but not sham stimulation. Since the effects of fixation offset in gap trials is mediated by subcortical mechanisms, mainly involving the inhibition of the superior colliculus, [Dorris and Munoz 1995], it is possible that the anodal tDCS influenced subcortical activity through cortico-subcortical connections with PPC and FEF.

### 4.3 Baseline performance

Of interest, an association between baseline performance and tDCS effects emerged, showing that “low performers”, namely participants with longer SRTs before stimulation showed a greater SRTs reduction after anodal tDCS, and this effect appearead to be area-, stimulus- and side-specific. Indeed, with respect to overlap trials, the baseline effect was greater after FEF stimulation, irrespective of direction of the saccades. Regarding the gap trials, instead, the association between baseline performance and tDCS effect was stronger after FEF and PPC stimulation, but only for leftward saccades. Overall, these results are consistent with previous evidence showing baseline-dependent effects of parietal tDCS on spatial attention [e.g., Learmonth et al. 2015], as well as of FEF tDCS on eye movements [Tseng et al. 2018]. Hence, it seems that the level of the baseline performance mediates side-and disengagement-specific effects of tDCS [e.g., Benwell 2015], but further research is needed to confirm this.

A final remark is concerned with some methodological aspects of tDCS. First, even though the electrode montage in the present study is quite standard, the computerized simulation in Figure 1 showed that the electrical current spread well beyond the target areas. This issue is known and could be addressed by employing electrodes and montages with higher spatial definition [Kuo et al. 2013; Bortoletto et al. 2016]. Finally, the adoption of an online tDCS approach may allow a better recruitment of structural and functional circuits of spatial attention and eye movements [Fertonani and Miniussi 2017].

## 5 CONCLUSION

The present study shows modulatory effects of anodal tDCS applied over PPC and FEF of the right hemisphere resulting in improvements of saccadic latency, regardless of the direction of the saccades. The effects seem to be more specific for gap trials, suggesting a possible involvement of subcortical mechanisms, but without definite evidence about the modulatory effects of tDCS on attentional disengagement. Importantly, the baseline level of performance influenced the tDCS effects.

## ACKNOWLEDGMENTS

This work was supported by the University of Milano-Bicocca (grant number: 2018-ATE-0317 to N.B.)

