## Supplemental Material for "Saccades, attentional orienting and disengagement: the effects of anodal tDCS over right posterior parietal cortex (PPC) and frontal eye field (FEF)"

### A APPENDICES

#### A.1 tDCS-related sensations

At the end of each tDCS session we administered a 7-item questionnaire, adapted from [Fertonani et al. 2015], to evaluate the potential adverse effects of tDCS. Participants were asked to report whether they felt 1) itching, 2) pain, 3) burning, 4) heat, 5) pinching, 6) metallic taste, or 7) fatigue, rating the intensity of their sensations using a 5-point scale (i.e., 0=absent, 1=Mild, 2=Moderate, 3=Considerable, 4=Strong). Moreover, they had to indicate when the feeling/discomfort began, when it stopped, where it was localized, and whether the it affected the performance. The reported sensations, their frequencies (out of 19 participants), and the most reported intensities can be found in table A.1.

**Table A.1: tDCS-related sensations, their frequency and the most reported intensity after frontal eye field (FEF), posterior parietal cortex (PPC), and sham stimulations**

| | FEF | | PPC | | sham | |
|---|---|---|---|---|---|---|
| **Item** | **N** | **Most reported** | **N** | **Most reported** | **N** | **Most reported** |
| Itching | 17 | Mild | 17 | Mild | 14 | Mild |
| Pain | 2 | Mild | 2 | Mild | 0 | |
| Burning | 12 | Mild | 10 | Mild | 8 | Mild |
| Heat | 10 | Mild | 8 | Mild | 8 | Mild |
| Pinching | 16 | Mild | 15 | Mild | 13 | Mild |
| Metallic Taste | 1 | Moderate | 0 | | 0 | |
| Fatigue | 5 | Moderate | 4 | Mild | 3 | Mild |

Overall, all participants localized the sensations on the head and these sensations were never rated as intense as “Considerable” or “Strong”.

For FEF stimulation, 18 out of 19 participants experienced these sensations at the beginning of the stimulation, whereas only one participant at the end. In the majority of participants, i.e., 12, sensations stopped quickly or in the middle of the stimulation, whereas for 7 participants these sensations stopped at the end of the stimulation. With respect to the influence on the task, only two participants reported that the feelings may have had a mild effect on the performance. For PPC stimulation, the sensations started at the beginning of the stimulation for all participants and stopped quickly or in the middle of the stimulation for most of them (16), whereas for 3 participants at the end of the stimulation. With respect to the influence on the task, only one participant reported that the feelings may have had a mild effect on the performance. For sham stimulation, one participant reported no feelings. 17 participants experienced these sensations at the beginning of the stimulation, 1 participant in the middle. For 16 out of 19 participants the sensations stopped quickly, whereas for 2 of them at the end of the stimulation. No participants reported that these sensations might have had an effect on the performance.

#### A.2 GLMM analyses of gap-overlap saccadic reaction times (SRTs)

Main effects and interactions of the gap-overlap SRTs analyses are reported in Table A.2.

**Table A.2: Main effects and interactions of gap-overlap SRTs analyses**

| Factors | $X^2$ | df | *p-value* |
|---|---|---|---|
| Stimulation | 9.13 | 2 | .010 |
| Timepoint | 41.3 | 1 | < .001 |
| Side | 5.31 | 1 | .021 |
| Trial Type | 3019.28 | 1 | < .001 |
| Stimulation * Timepoint | 7.71 | 2 | .021 |
| Stimulation * Side | 1.7 | 2 | .427 |
| Timepoint * Side | .03 | 1 | .859 |
| Stimulation * Trial Type | 1.75 | 2 | .418 |
| Timepoint * Trial Type | 4.28 | 1 | .039 |
| Side * Trial Type | 13.48 | 1 | < .001 |
| Stimulation * Timepoint * Side | .98 | 2 | .612 |
| Stimulation * Timepoint * Trial Type | 1.26 | 2 | .533 |
| Stimulation * Side * Trial Type | .1835 | 2 | .204 |
| Timepoint * Side * Trial Type | .46 | 1 | .498 |
| Stimulation * Timepoint * Side * Trial Type | .09 | 2 | .957 |

Regarding the main effect of *Stimulation*, SRTs associated to PPC (*M*=205 ms; *SE*=4.35 ms) were slower (*z*=-3.03; *p*=.007) than those associated to FEF (*M*=201 ms; *SE*=4.32 ms). No difference emerged between sham SRTs (*M*=203 ms; *SE*=4.32 ms) and FEF (*z*=-1.61; *p*=.325) or PPC (*z*=1.42; *p*=.468). With respect to *Timepoint*, SRTs were significantly faster after the stimulation (*M*=200 ms; *SE*=4.3 ms) than before stimulation (*M*=206 ms; *SE*=4.29 ms). As for the *Side*, overall SRTs for right saccades (*M*=202 ms; *SE*=4.29 ms) were slightly faster than for left saccades (*M*=204 ms; *SE*=4.3 ms). Concerning the interaction *Stimulation*Timepoint*, SRTs were faster after both FEF stimulation (FEF pre: *M*=204 ms; *SE*=4.4 ms; FEF post: *M*=198 ms; *SE*=4.41 ms; *p*=.003), and PPC tDCS (PPC pre: *M*=210 ms; *SE*=4.42 ms; PPC post: *M*=200 ms; *SE*=4.45 ms; *p*<.001), but not after sham stimulation (sham pre: *M*=205 ms; *SE*=4.41 ms; sham post: *M*=201 ms; *SE*=4.41 ms; *p*=.1). The interaction *Timepoint*Trial Type* revealed that the decrease of SRTs was stronger for gap trials (pre: *M*=179 ms; *SE*=4.34 ms; post: *M*=170 ms; *SE*=4.34 ms; *p*<.001) than overlap trials (pre: *M*=234 ms; *SE*=4.37 ms; post: *M*=229 ms; *SE*=4.39 ms; *p*=.036). Finally, the interaction of *Side*Trial Type* showed that, in overlap trials, SRTs for right saccades (*M*=228 ms; *SE*=4.37 ms) were significantly faster as compared to left saccades (*M*=235 ms; *SE*=4.39 ms; *p*=.001), whereas no significant difference was found for the gap trials (left saccades: *M*=174 ms; *SE*=4.34 ms; right saccades: *M*=175 ms; *SE*=4.33 ms; *p*=1).

##### A.3 Analyses of the Gap Effect (GE)

Main effects and interactions of the GE analyses are reported in Table A.3

**Table A.3: Main effects and interactions of GE analyses**

| Factors | F | df | *p-value* |
|---|---|---|---|
| Stimulation | .36 | 2, 198 | .697 |
| Timepoint | 3.56 | 1, 198 | .061 |
| Side | 5.17 | 1, 198 | .024 |
| Stimulation * Timepoint | .06 | 2, 198 | .938 |
| Stimulation * Side | .9 | 2, 198 | .408 |
| Timepoint * Side | .23 | 1, 198 | .635 |
| Stimulation * Timepoint * Side | .14 | 2, 198 | .868 |

##### A.4 Explorative analyses for gap and overlap trials

Non-aggregated raw SRTs for both gap and overlap conditions were heavily right-skewed. Thereupon, for both gap and overlap trials, we tested a three-way *Simulation* (FEF, PPC, and sham)**Timepoint* (pre- and post-stimulation)**Side* (left and right saccades) term by employing Gamma GLMMs that regarded subjects as clusters. A random *Stimulation***Time point* term was entered but yielded singularities; therefore, a random intercept only was fitted in both models. Deviation coding was adopted for factors. Bonferroni-corrected post-hoc comparisons were also performed.

As for the analyses of gap trials a trend was observed for *Stimulation* ($\chi^2(2)=5.34$; $p=.069$). Moreover, the effect of *Timepoint* was significant ($\chi^2(1)=53.49$.; $p<.001$), i.e., SRTs were overall significantly higher prior to (*M*=178 ms; *SE*=4.94 ms) compared to post-stimulation (*M*=169 ms; *SE*=4.94 ms). Importantly, a significant *Stimulation*Time point* interaction ($\chi^2(2)=10$; $p=.007$) showed that SRTs significantly decreased post-stimulation for both FEF (pre: *M*=176; *SE*=5.09; post: *M*=167; *SE*=5.08; $p=.001$) and PPC (pre: *M*=182 ms; *SE*=5.09 ms.; post: *M*=168 ms; *SE*=5.08 ms; $p<.001$), whereas not for Sham tDCS (pre: *M*=176 ms; *SE*=5.10.; post: *M*=172 ms; *SE*=5.07; $p=.757$). Notably, we did not observe significant baseline differences, i.e., prior to stimulation, neither between FEF and PPC ($p=.182$), between FEF and sham ($p=1$), nor between PPC and sham tDCS ($p=.252$).

Analyses of the overlap trials revealed a trend for the main effect of *Stimulation* ($\chi^2(2)=5.95$; $p=.051$). Moreover, concerning the effect of *Timepoint*, SRTs were overall significantly higher ($\chi^2(1)=7.31$; $p=.007$) before tDCS (*M*=236 ms; *SE*=5.17 ms) as compared to after stimulation (*M*=231 ms; *SE*=5.15 ms). Furthermore, the main effect of *Side* ($\chi^2(1)=12.32$; $p=<.001$) showed that rightward saccades (*M*=231; *SE*=5.15) were faster than leftward ones (*M*=236 ms *SE*=5.17 ms). No significant interactions were detected. Result of gap analyses are reported in Table A.4 and depicted in Figure A.1, whereas results of overlap analyses can be found in table A.5 and Figure A.1

**Table A.4: Main effects and interactions for gap SRTs**

| **Factors** | **X²** | **df** | ***p-value*** |
|---|---|---|---|
| Stimulation | 5.34 | 2 | .069 |
| Timepoint | 53.5 | 1 | < .001 |
| Side | 1.02 | 1 | .312 |
| Stimulation * Timepoint | 10 | 2 | .007 |
| Stimulation * Side | .24 | 2 | .885 |
| Timepoint * Side | .4 | 1 | .528 |
| Stimulation * Timepoint * Side | .85 | 2 | .654 |

**Table A.5: Main effects and interactions for overlap SRTs**

| **Factors** | **X²** | **df** | ***p-value*** |
|---|---|---|---|
| Stimulation | 5.95 | 2 | .051 |
| Timepoint | 7.31 | 1 | .007 |
| Side | 12.32 | 1 | < .001 |
| Stimulation * Timepoint | 1.41 | 2 | .495 |
| Stimulation * Side | 3.58 | 2 | .167 |
| Timepoint * Side | .17 | 1 | .683 |
| Stimulation * Timepoint * Side | .75 | 2 | .688 |

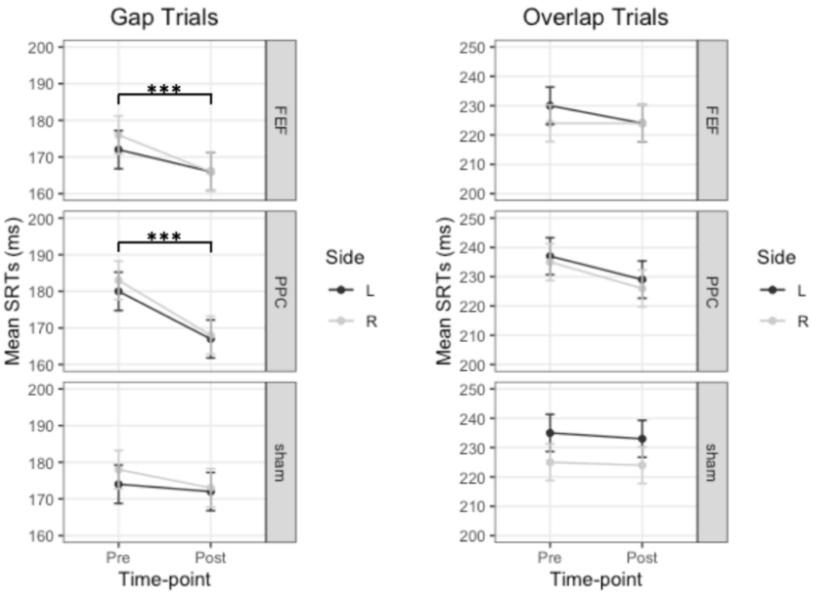


**Figure A.1: Main results of gap and overlap saccadic reaction times (SRTs) according to *Stimulation*, *Timepoint*, and *Side* of saccade. The significant *Stimulation*Timepoint* for gap trials is also depicted. Values represent estimated marginal means and the standard error of the mean. FEF= frontal eye field; PPC= posterior parietal cortex; L= left; R= right. ***= *p*<.001.**
